# Utilizing Q-Learning to Generate 3D Vascular Networks for Bioprinting Bone

**DOI:** 10.1101/2020.10.08.331611

**Authors:** Ashkan Sedigh, Jacob E. Tulipan, Michael R. Rivlin, Ryan E. Tomlinson

## Abstract

Bioprinting is an emerging tissue engineering method used to generate cell-laden scaffolds with high spatial resolution. Bioprinted vascularized bone grafts are a potential application of this technology that would meet a critical clinical need, since current approaches to volumetric bone repair have significant limitations. However, generation of vascular networks suitable for bioprinting is challenging. Here, we propose a novel Q-learning approach to quickly generate 3D vascular networks within patient-specific bone geometry that are optimized for bioprinting. First, the inlet and outlet locations are specified and the scenario is modeled using a grid world for initial agent training. Next, the path planned in the grid world environment is converted to a Bezier curve, which is then used to generate the final 3D vascularized bone model. The vessels generated using this procedure have minimal tortuosity, which increases the likelihood of successful bioprinting. Furthermore, the ability to specify inlet and outlet position is necessary for both surgical feasibility as well as generation of more complex vascular networks. In total, this study demonstrates the reliability of our reinforcement learning method for automated generation of 3D vascular networks within patient-specific geometry that can be used for bioprinting vascularized bone grafts.

## I. Introduction

Regenerative medicine is an emerging field that seeks to develop methods to regrow or replace damaged, diseased, or missing tissues with synthesized tissue that restores normal function [1][2]. The development of a regenerative medicine approach to generate new bone and cartilage for treatment of degenerative joint diseases has been an active research area for many years [3]. However, the bone and/or cartilage constructs generated using standard tissue engineering strategies lack the spatial complexity of native tissue. In this regard, bioprinting, which utilizes 3D printing technology to generate tissue using materials containing viable cells, may be a solution for generating patient-specific tissue for bone grafting [4].

Bioprinting is a growing field that is expected to significantly impact clinical practice by enabling new regenerative medicine approaches. In general, bioprinted constructs are generated by sequentially printing thin layers of specific materials, such as hydrogels, collagen, and bioceramics, that are laden with cells. The layer geometry is stored in a G-code file that the bioprinter translates to extrude the particular material. With the potential to produce a specific 3D shape containing cells at high resolution, 3D bioprinting has become a popular biofabrication method for researchers [5]. In particular, bioprinting a vascularized bone tissue construct would be a significant improvement over current efforts [6][7] and would directly meet a pressing clinical need.

In particular, strategies to repair large bone defects in humans in scenarios such as neoplasm, trauma, reconstruction, and infection are limited [8]. Osteoconductive bone substitutes can be used to provide a scaffold for mineralization by native osteoblasts but are limited by the slow rate and limited reach of bony ingrowth, technical difficulties shaping the construct, and concerns about structural strength [9]. Similarly, bone allograft is also limited by the rate of host bone ingrowth and has the added complications of donor availability and attritional weakening of the allograft [10]. Although autograft bone contains living cells able to produce new bone, it cannot be used to treat large defects due to diffusion limiting the ability of cells in the center of a large graft mass to obtain nutrients and remove waste products, ultimately resulting in fatigue failure [11]. Vascularized bone grafts, which utilize the native vascularity of a bone graft to accomplish nutrient and waste exchange, were introduced to address this major limitation [12]. Unfortunately, vascularized bone donor sites are limited in number, size, and contour. Furthermore, harvesting these grafts results in added patient morbidity, and their implantation requires considerable technical skill [13]. In total, there are many clinical scenarios of volumetric bone loss lacking a suitable method for treatment.

Since the skeletal vascular network is critical to native bone mineralization and graft survival [14], bioprinted bone intended for the treatment of large defects must be effectively vascularized. As a result, this requirement necessitates the design of a 3D vascular network within the bioprinted bone. We have developed a novel method to implement a vascular network using patient-specific geometry at the desired vascular density with customizable inlet and outlet positions by optimizing tortuosity using Reinforcement Learning (RL). This paper introduces the implemented learning method and shows the mathematical convergence and validation for the learning method (termed Q-Learning). This method presents our 3D-to-2D projection, agent training, and a proper learning environment called the grid world path planning. Bezier curve approximation and 2D-to-3D methods are described as the final steps to implement the imported geometry’s computed vessels.

## II. Q-Learning

Reinforcement Learning (RL) is a semi-supervised learning method that solves a task by trial and error by acting within an environment and calculating the feedback rewards for each taken action in order to maximize the accumulated reward [15], [16]. For each time step in which the agent takes an action, the environment transitions to a new state. The environment feedback is less informative than supervised learning and more informative than unsupervised learning since agents in unsupervised learning must discover the world without any explicit feedback [17].

RL contains Monte-Carlo learning, temporal difference, and dynamic programming learning. Q-learning and State-Action-Reward-State-Action (SARSA) are the two algorithms of temporal difference learning [18]. Single-agent RL algorithms are divided into both model-free and model-based methods. Model-based methods include dynamic-programming; on the other hand, model-free methods are based on an online estimation [17].

Markov Decision Process (MDP) describe the agent environment by the following definition.

### Definition 1

A Markov decision process is defined as a tuple *M* = (*X, A, p, r*) where *X* is the countable, finite and continuous state space, *A* is the finite, continuous, and countable action space. For the dynamic environments, the transition probability is *p*(*y*|*x, a*) for any *x ϵ X, y* ∈ *X*, and *a ϵ A*. Equation (1) is the probability of observing a next state *y* when an agent take action *a* in the state space *x*.

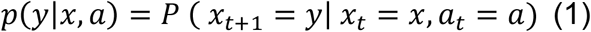

### Definition 2

Policy is a decision rule *π*_*t*_ is a state to action mapping at the time *t* ∈ ℕ which is define as equation (2). In a Markovian process, policy is the sequence of decision rules *π* = (*π, π, π*, …).

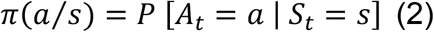

The agent’s goal is to maximize the expected discounted return at each step time *t*, as shown in equation (3):

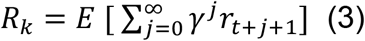

In equation (3), γ ∈ [0,1) is the discount factor, which is considered as the uncertainty regarding the received rewards in the future. Small *γ* looking for short-term rewards and values close to 1 look for the long-term rewards, which result in the exploration versus exploitation criteria. *R*_*t*_ represents the agent reward accumulated in the long process. According to the equation (3), in order to calculate the state value for infinite time with a discount factor, equation (4) is used:

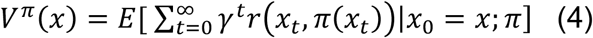

### Definition 3

the state value function or *Q*-function for any policy *π, Q*^*π*^: *X* × *A* → ℝ is defined as equation (5):

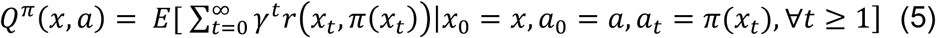

And the optimal *Q*-function describes as *Q*\*(*x, a*) = *max*_*π*_*Q*^*π*^(*x, a*) as we deduce that the optimal policy is *π*\*(*x*) = arg *max*_*a*∈*A*_ *Q*\*(*x, a*). The Bellman optimality equation defined as equation (6):

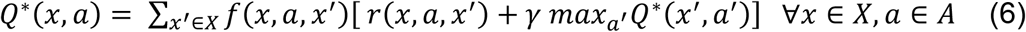

Equation (6) states that the current value by taking action *a* in state space *x* is the expected immediate reward plus the optimal policy (discounted) from the future states *x. x*^′^ *indicates the next state in the environment*. In the RL algorithm, the action goal is to maximize the return by choosing actions with *epsilon greedy* and the optimal *Q*\*.

Q-learning is an online estimation with a model-free learning method [19], [20]. It turns to a learning algorithm by putting the equation (6) into an iterative loop. The *Q*\* is estimated using samples of equation (6). The sample batch is computed in the environment by reward *r*_*t*+1_, and the states of *x*_*t*_, *x*_*t*+1_:

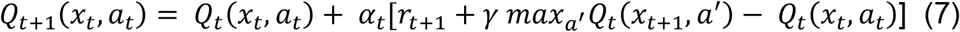

The equation (7) does not have any information regarding the transition probability and reward functions; therefore, Q-learning is a model-free algorithm. The parameter *α*_*t*_ ∈ (0,1] is the time-varying learning rate that specifies how far steps can be taken to determine the value of the batch sample (target) γ *max*_*a*_′ *Q*_*t*_(*x*_*t*+1_, *a*^′^) − *Q*_*t*_(*x*_*t*_, *a*_*t*_). The convergence of the equation (7) has been considered and mathematically proven under the following conditions[17], [21]:

- Q-learning updated values must be stored for each state action *Q*_*t*_(*x*_*t*_, *a*_*t*_)
- The series of time-varying learning rate for each state action (*x*_*t*_, *a*_*t*_) sums infinity, but the sum of its square should be finite[22]:

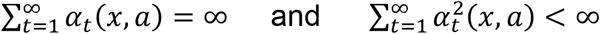
- The agent should explore the environment in all states with nonzero probability.

In order to guarantee the third condition for the agent, a greedy policy is used. In this condition, at each step, the agent chooses a random action with probability of *ε* ∈ (0,1), and the greedy action with the probability (1 − *ε*). This *ε*-greedy technique is used to explore the environment rather than exploit in one action[23]. Another method, which is the Boltzmann exploration strategy, can be used to find the action probability by purely random action selection [24].

The difference between SARSA and Q-Learning algorithm is the Q-function update as indicated equation (8) vs (7). In the SARSA algorithm, it computes the difference between *Q*_*t*_(*x*_*t*_, *a*_*t*_) and the weighted sum of the average action value and the maximum Q Value. In the SARSA algorithm, the target policy is always same as the behavior policy [25]:

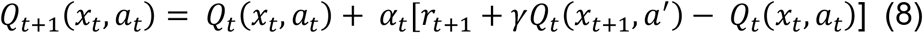

## III. Methodology

The implementation of Q-learning for the generation of a 3D vascularization model based on the raw image data involves the following steps: In the first step, cross-sectional 2D images, such as those generated by CT or MRI medical imaging, will be converted into the 3D model to specify the inlet and outlet position of the vascular network. In the next step, the required vascularization density and number of the vessels will be specified and the 3D model will be sliced into 2D planes. In order to simulate the Q-learning algorithm to find the solution, which is the least tortuosity and least overall distance to the outer shells, the 2D slice is converted to a 2D grid plane containing both the inlet and outlet position. Tortuosity index is the ratio of the total length and preferential tortuous fluid pathways.

Q-learning solution, which is a planned path, is converted to a 2D Bezier curve and a 3D shape with the specified diameter. This 3D vascularized model is then implemented by subtraction from the initial 3D solid bone model. This results in a 3D vascularized bone model that can be 3D printed or used for *in silico* simulation.

### A. 3D Model Reconstruction and Slicing

Medical imaging techniques, such as CT or MRI, are in widespread use for visualizing musculoskeletal anatomy and pathologies. Therefore, these data can reasonably be used to extract patient-specific 3D geometry for bioprinting [26]. One way to reconstruct the 3D model is to detect the contour in each cross-sectional image, then construct the mixed layers in a triangular STL model [27][28].

We used a human scaphoid bone for implementation of the Q-learning algorithm, which is a non-convex shape. Figure 1 shows the generated mesh of human scaphoid from CT scan data. The inlet and outlet position on the vessel are essential for optimal clinical use. As indicated in the algorithm (1), the inlet and outlet position coordinates are saved for the further use in the algorithm (2). The generated 3D mesh is sliced to convert the 3D model into 2D slices. The algorithm computes each slice area and chooses the maximum size as the target plane for implementing the vascularization network. This plane is generated by the specific Z position passing from the inlet and outlet pairs *(x*_*i*_, *y*_*i*_, *z*_*i*_) and *(x*_*o*_, *y*_*o*_, *z*_*o*_). The next step is to generate the vascularization network in the newly generated 2D plane.

**Figure 1.**
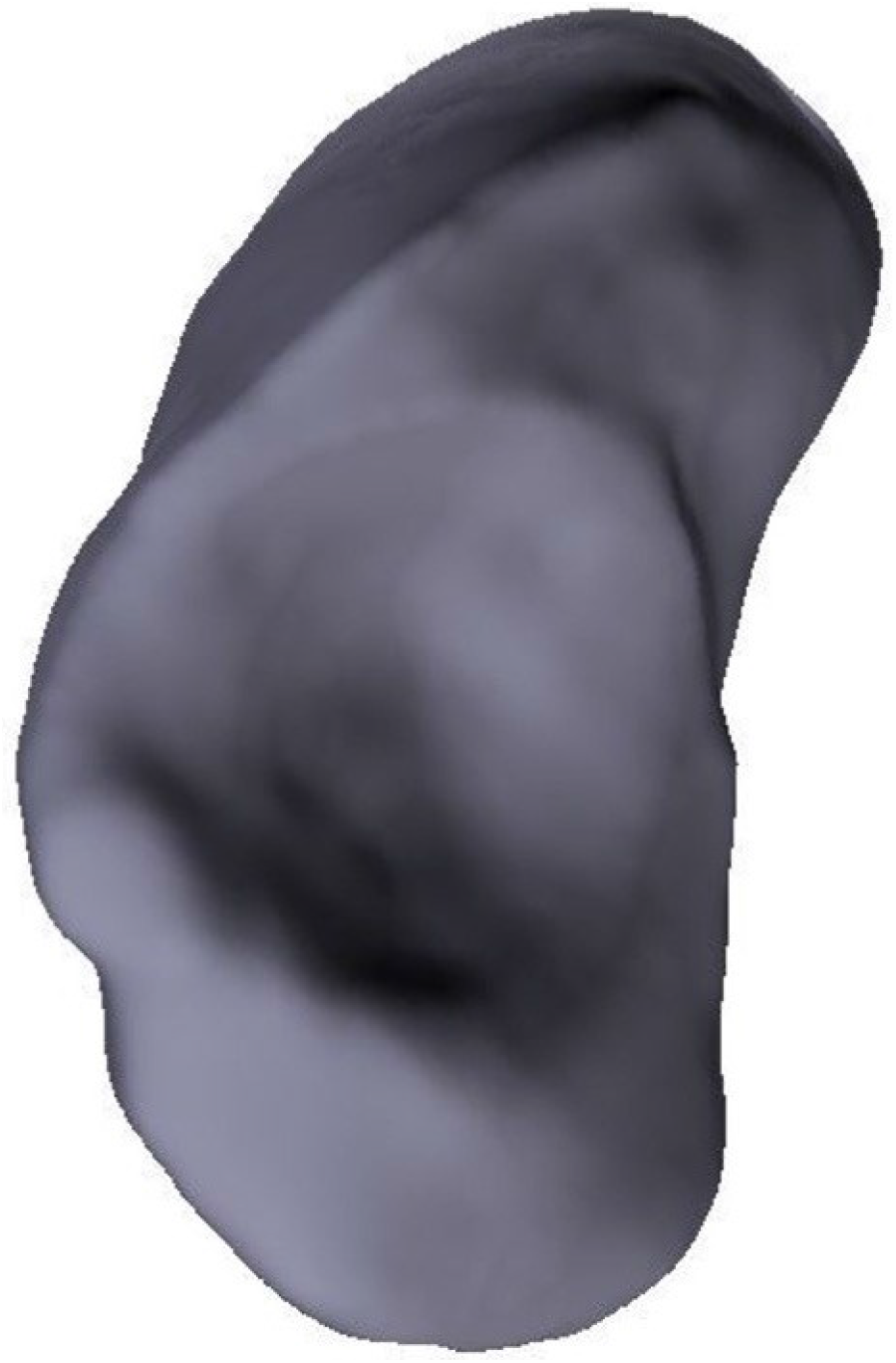
Scaphoid 3D Model. Reconstruction of the scaphoid bone from imaging data illustrates its non-convex surface.

#### Algorithm 1 3D Reconstruction and Slicing

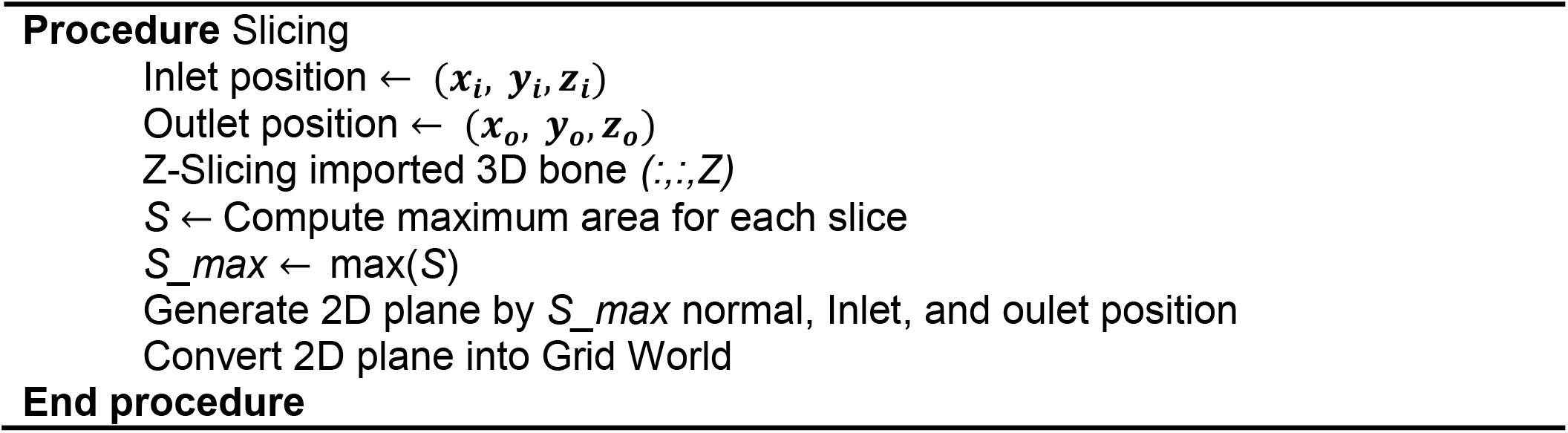

### B. Q-learning

Path planning vascularization in a 2D plane has several constraints. The algorithm should consider both the tortuosity index and the coverage area by measuring curvature distance to the model’s outer shells. Therefore, it is required to look for an algorithm that can find a 2D space solution with numerous possibilities. Algorithm (2) is the general workflow for this aim.

2D grid plane from part A is generated by choosing the scaphoid slice’s maximum length and width. This 2D slice is converted to the grid world as the RL algorithm environment. In the next step, the Q-learning algorithm is set up to solve this problem by maximizing reward. The policy for each position shows the path and agent decision to move in this plane. This simulation is performed with MATLAB® R2020a and Reinforcement Learning Toolbox.

RL grid world problem consists of three main components:

- **Agent:** in this scenario, as illustrated in Figure 2A, agent is the vessel that completes a path which will be used for vessel modeling. The agent does action *a* and the environment, which is a 2D plane, returns the *r*_*t*+1_ and *x*_*t*+1_, which is the next state. Agent training parameters, episode information, and average results are shown in Table 1.
- **Goal:** this scenario aims to start from the inlet position and finish it at the outlet position.
- **Obstacles:** These obstacles are in black, as shown in Figure 2B. Agents should avoid these obstacles out of path planning areas or distance to the outer contour. The agent will receive a negative reward signal by passing over obstacles.

This algorithm aims to train an agent to reach the goal of avoiding obstacles in the grid world so that the accumulated rewards by movements get maximized. To accomplish this aim, the agent has to discover the world and learn the environment’s dynamics. A proper value for each environment section as movement, goal, and obstacles is defined. Colliding obstacles or defined boundaries, a highly negative reward signal is given to the agent.

**Table 1.** Training Parameters

| Agent Training Parameters | Value |
| --- | --- |
| Maximum Steps Per Episode | 50 |
| Epsilon | 0.4 |
| Stop Training Value | 1000 episodes |
| Maximum Number of Episodes | 1000 |
| Elapsed Time | 108 seconds |
| Episode Steps | 13 |
| Episode Reward | -50 |
| Episode Q0 | -47 |
| Total Number of steps | 22966 |
| Average Reward | -57 |
| Average Steps | 14.4 |

**Figure 2.**
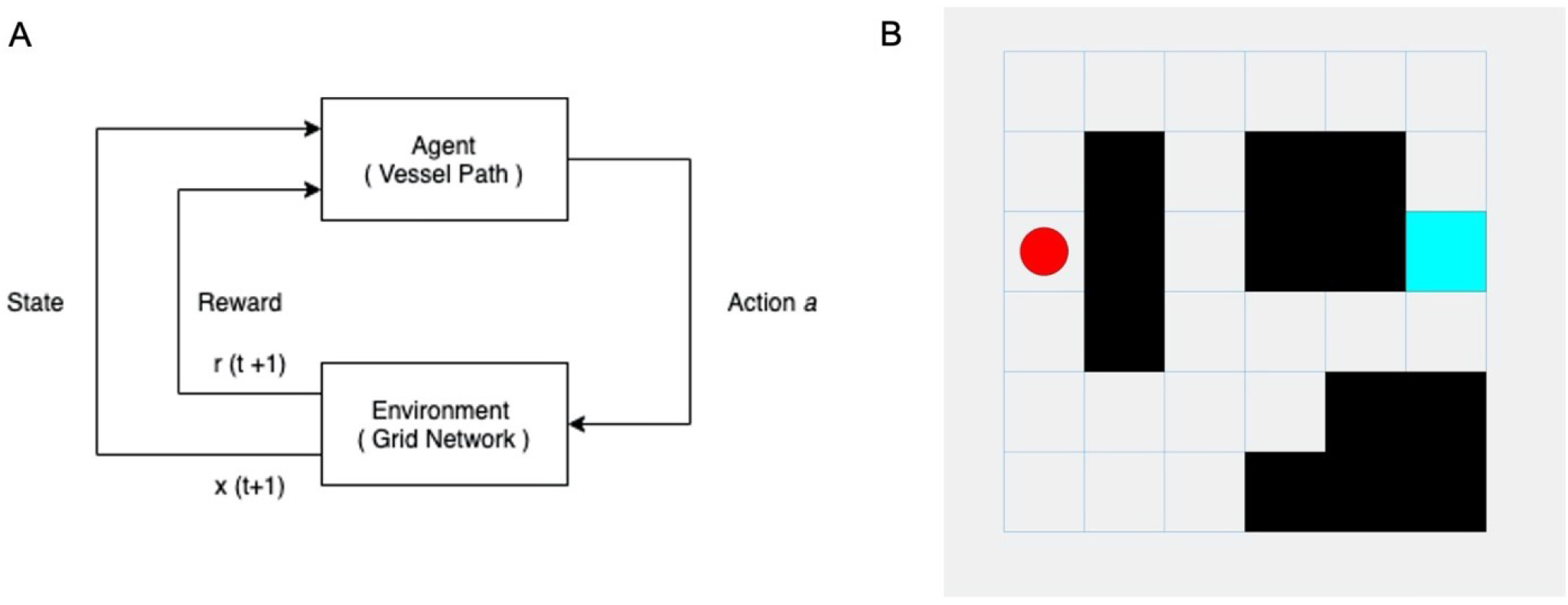
Reinforcement Learning Environment. A) The Reinforcement Learning workflow in which the agent (vessel) takes action *a* in the grid world environment. The environment returns *r*_*t+1*_ and the next state *x*_*t+1*_. B) The grid world used to train the agent to find the path with least tortuosity between the inlet (red) and outlet (blue), where obstacles (defined avascular areas or boundaries) are black.

#### Algorithm 2 Q-Learning

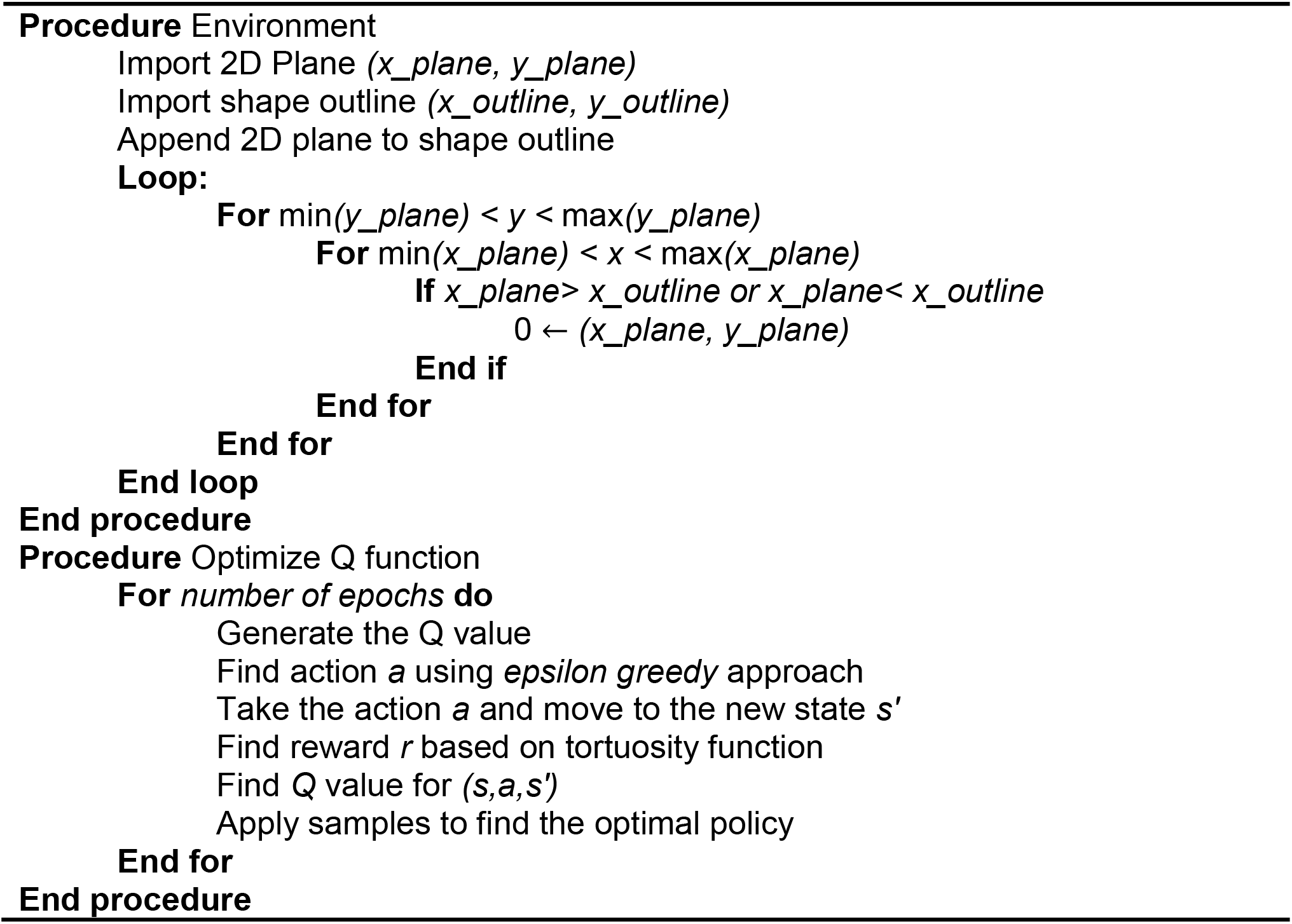

### C. Bezier Curve Approximation and 3D Rendering

Bezier curves were introduced by Paul de Casteljau in early 1960s [29].This algorithm is based on the interpolation between the pair of control points. A Bezier curve with degree of *n* needs *n+1* control point *b*_*i*_ ∈ *R*^*d*^, *i* = 0, 1, …, *n, t* ∈ *R*:

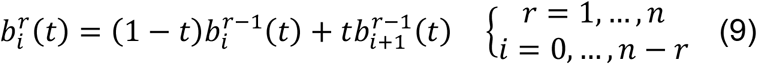

And 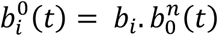 is the point with parameter *t* on the Bezier curve *b*^*n*^. The polygon P which is calculated by *b*_0_, …, *b*_*n*_is called the *Bezier polygon* of the curve *b*^*n*^. *b*_*i*_ are called control points [30].

Agent path planning solution as pairs of (*x*_*path,n*_, *y*_*path,n*_) are considered as the control points *b*_0_, …, *b*_*n*_. Therefore, the planned path turns to a 2D Bezier curve as Figure 3. For making a 3D path with a desired diameter, a Python script on Blender [31] is programmed to convert the 2D path into 3D model. The desired diameter is considered as a relation of the difference in pressure (Δ*P*), the viscosity of the fluid (*μ*), and the vessel length (*L)* [14]:

**Figure 3.**
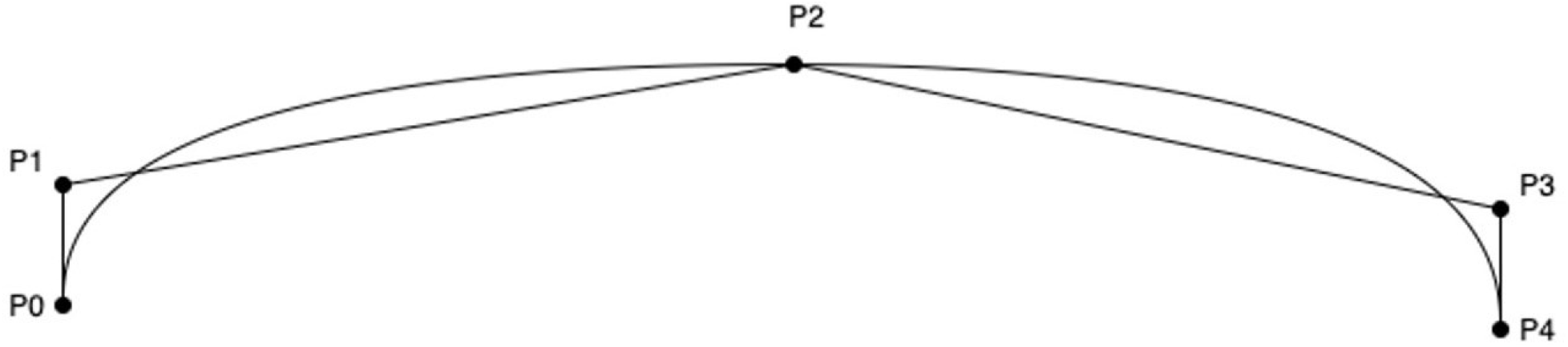
Bezier Curve. A vessel as defined by the Bezier Curve calculated from the four control points [*p1,p2,p3,p4*].

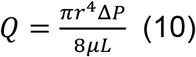

The final 3D Bezier curve will be implemented starting from the inlet position to the outlet position as pairs of *(x*_*i*_, *y*_*i*_, *z*_*i*_*) (x*_*o*_, *y*_*o*_, *z*_*o*_).

## IV. Results

Different scenarios for 3D vascularization networks based on the grid world path planning with varying constraints of reward have been tested. Table 2 shows the different scenarios including number of episodes, episode Q0, average reward, and tortuosity index as a result. Figure 4A-E illustrates the different planned path based on the various reward function constraints. To validate the Q-Learning algorithm’s training status using the planned path the training status for each Q0 episode, average reward, and episode reward have been plotted Figure 4F. Here, the final episode’s value is converted to the desired reward value, which indicates the tortuosity index level as low, medium, or high. Algorithm picks the result which is converged and has the minimum value of tortuosity index. In this example the path planned with 1000 number of episodes is the solution for generating a vessel since it has the least tortuosity index and the plot is converged.

**Table 2.** Comparison of path planning in a grid world

| Maximum number of Episodes | Average Rewards | Tortuosity Index | Episode Q0 | Converged |
| --- | --- | --- | --- | --- |
| 200 | -78.8 | 1.4 | -49.9 | No |
| 400 | -60 | 1.3 | -56 | Yes |
| 700 | -77 | 1.3 | -56 | No |
| 1000 | -57 | 1.16 | -47 | Yes |
| 1500 | -114 | 1.5 | -65 | No |

**Figure 4.**
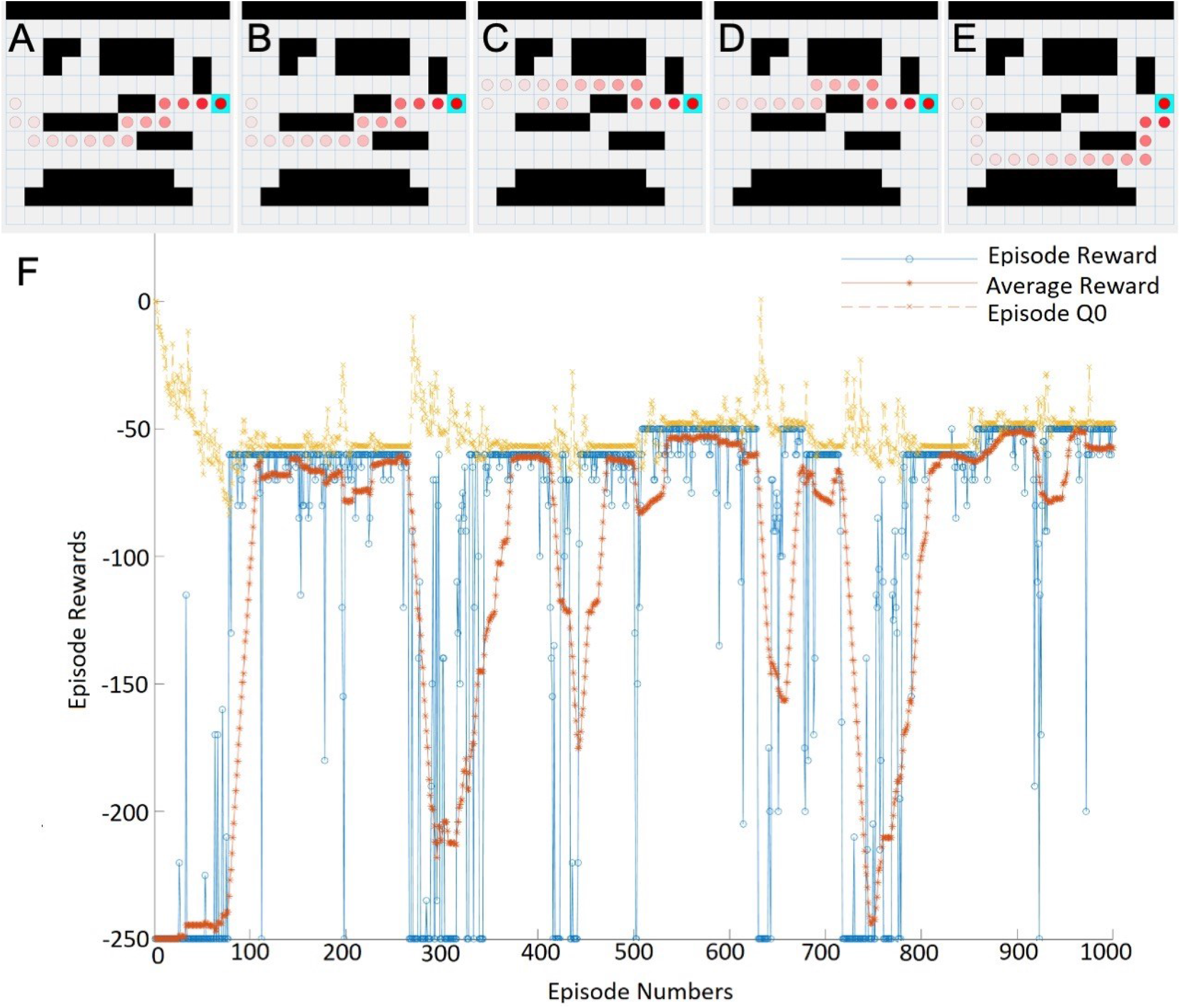
Path Planning in grid world and RL Training result. A-E) Path planned for vessel transiting from the inlet to the outlet in the grid world following A) 200, B) 400, C) 700, D) 1000, and E) 1500 episodes. F) Results of Q-learning algorithm for 1000 episodes.

Average reward reinforcement learning algorithms convergence follows the idea from a kind of Tauberian theorem; if the discount rate converges to one, it is converged to the average reward value [32]. The fact that episode reward and average reward are converged to the same value indicates the Q-learning algorithm convergence and validation of the training algorithm.

3D vascularized scaphoid model is generated with Blender python scripts and Bezier curve approximation algorithm, as shown in Figure 5A. Using this algorithm, it is possible to generate a second vessel to increase vascular coverage in a large construct. To do so, the inlet and outlet of the second vessel is located at the main vessel with a larger diameter before and after the smaller vessel inlets and outlets. This scheme results in a more complex vascular network that remains compatible with 3D bioprinting (Figure 5B).

**Figure 5.**
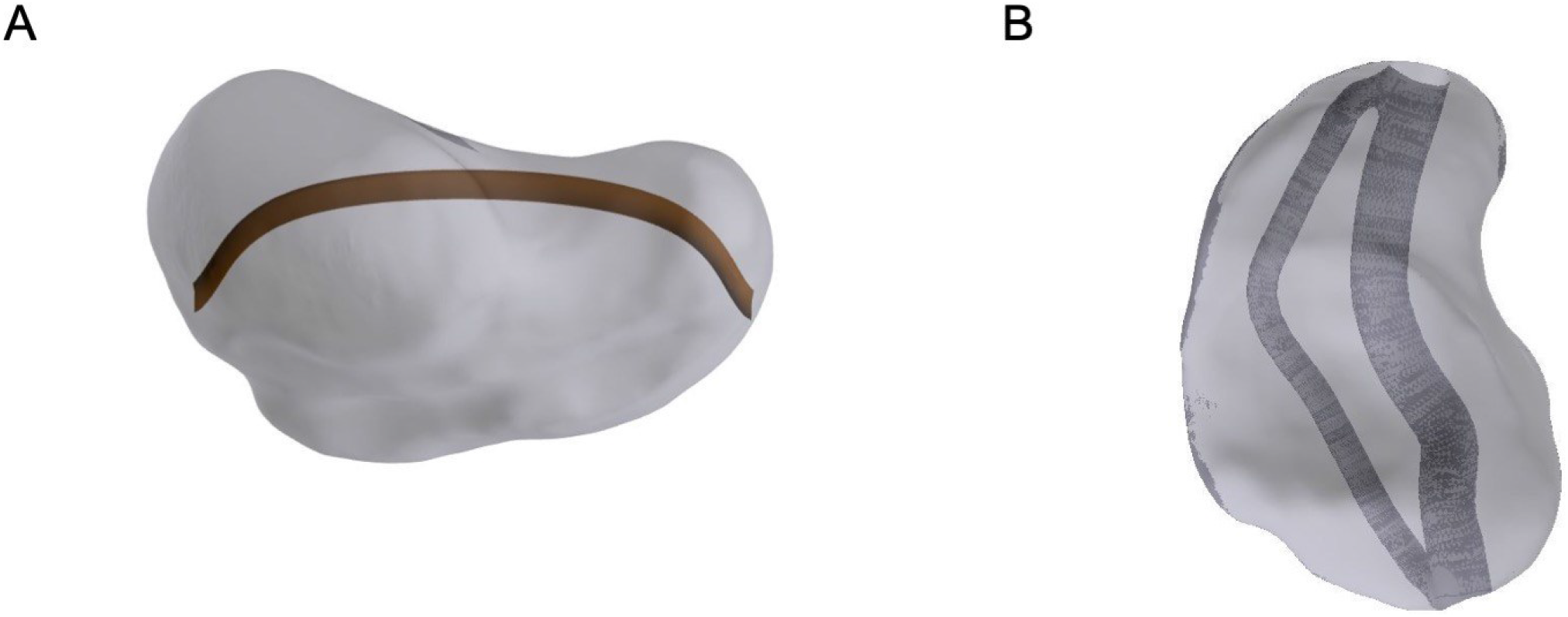
Rendering of vessel generated by Q-learning. A) Path planned for the scaphoid geometry by Q-learning has been converted to a Bezier curve and implemented in 3D. B) Path planned for a second vessel within vascularized scaphoid by locating the inlet and outlet on the first vessel.

## V. Conclusion

A model-based RL algorithm was used to generate a vascular network in patient-specific geometry with the specified inlet and outlet position, vascularization density, and tortuosity index. This model-based RL algorithm has an efficient training method without considerable computation time. Furthermore, the ε-greedy Q-learning approach is a method requiring minimal computational resources for training agents in this path planning problem. With the slicing method, the grid world environment for the RL agent is extracted for the simulation. The data from this simulation was used to find the optimal policy regarding the minimum tortuosity and maximum area coverage. Finally, the planned path was converted to the Bezier curve with an approximation and then converted into a 3D model, which was then implemented in the initial 3D bone model.

Here, the generated 3D model was 3D printed to validate the geometry and vessel functionality after optimization. We note that this model simulates a single agent in the grid world, and thus limits the grid world’s multi-vessel generation flexibility. In future studies, it may be required to implement a multi-agent path planning algorithm to optimize the number of vessels required for a more complex model. One such example would be the implementation of multiple independent vascular networks in the same bone construct.

Current techniques in bone grafting, and complex engineered tissues generally, are limited in size by the metabolic demands of living cells that exceed the limit of diffusion. As a result, semi-automated machine-learning generation of vascular networks provides a critical functionality for advancing bioprinted bone constructs. In turn, this study moves the field one step closer to routine clinical use of large volume, patient-specific bioprinted tissue grafts.

## Author Contributions

Conceptualization (AS, JET, MRR, RET), Investigation (AS, JET, MRR, RET), Writing (AS, JET, MRR, RET), Funding (MRR, RET).

## Acknowledgements

Our research is supported by the National Institute of Arthritis and Musculoskeletal and Skin Diseases and the National Institute of Dental and Craniofacial Research of the National Institutes of Health under award numbers AR074953 (RET) and DE028397 (RET). The content is solely the responsibility of the authors and does not necessarily represent the official views of the funding bodies.

